# Eye pupil signals information gain

**DOI:** 10.1101/693838

**Authors:** Alexandre Zénon

## Abstract

In conditions of constant illumination, the eye pupil diameter indexes the modulation of arousal state and responds to a large breadth of cognitive processes, including mental effort, attention, surprise, decision processes, decision biases, value beliefs, uncertainty, volatility, exploitation/exploration trade-off or learning rate. Here, I propose an information theoretic framework that has the potential to explain the ensemble of these findings as reflecting pupillary response to information processing. In short, updates of brain internal model, quantified formally as the Kullback-Leibler (KL) divergence between prior and posterior beliefs, are the common denominator to all these instances of pupillary dilation to cognition. I show that stimulus presentation leads to pupillary response that is proportional to the amount of information the stimulus carries about itself and to the quantity of information it provides about other task variables. In the context of decision making, pupil dilation in relation to uncertainty is explained by the wandering of the evidence accumulation process, leading to large summed KL divergences. Finally, pupillary response to mental effort and variations in tonic pupil size are also formalized in terms of information theory. On the basis of this framework, I compare pupillary data from past studies to simple information theoretic simulations of task designs and show good correspondance with data across studies. The present framework has the potential to unify the large set of results reported on pupillary dilation to cognition and to provide a theory to guide future research.

## Cognitive pupillary response

Beside the well-known response of pupillary muscles to light, allowing to narrow the range of light intensity reaching the retina and optimizing its information capacity [1], pupil size varies also as a function of a wealth of cognitive phenomena, including mental effort [2, 3, 4, 5], surprise [6, 7, 8, 9, 10, 11, 12, 13, 14, 15], emotion [16], decision processes [17, 18, 19, 20], decision biases [21, 19, 22], value beliefs [23, 24, 25], volatility (unexpected uncertainty; [26, 27, 28, 10], exploitation/exploration trade-off [29, 30], attention [31, 32, 33, 34, 35, 36], uncertainty [37, 19, 38, 12, 21, 23, 25], confidence [39], response to reward [40], learning rate [41, 10, 42, 43, 41], neural gain [44, 10, 36, 45] or urgency [46]. These variations in diameter follow coherent changes in neural activity throughout cortex, regulated by neuromediators, and referred to as *arousal* [47, 48, 49, 50]. The present work is based on the strong hypothesis that the ensemble of phenomena that trigger changes in pupil-linked arousal all depend on a basic underlying information theoretic process: the update of brain internal models. We will review a large breadth of findings from the literature and will reinterpret them under the light of that framework.

## Surprise and self-information

One of the first cognitive variables that was shown to influence pupillary responses is surprise, defined in information theory as the negative logarithm of the probability of an event. This quantity is also called self-information, because it measures how much information is gained when observing an event. Pupil size has been shown to respond vigorously and robustly to surprise, with dilation in response to events in inverse proportion to their frequency of occurrence in a trial [51, 14, 52]. Pupil also responds to stimulus disappearance, in inverse proportion to how likely the stimulus is to disappear at that given time [9]. Along the same line, pupillary dilation has been reported in relation to the probability of a reward outcome, independently of its sign (i.e. responses are equivalent for losses and rewards; Van Slooten et al. [25], Lavín et al. [6], Satterthwaite et al. [37]) or even to the occurrence of errors, as a function of their likelihood [53]. When events have probability distributions defined along continuous feature spaces (e.g. position, number line), pupil also responds in inverse proportion to the probability density of occurrence of that feature [13, 10]. When event occurrences depend on past trial history, pupil responses reflect surprise taking account of that history [12, 10]. Despite this apparent consistency of findings, no attempts have been made so far to assess whether the relationship between pupil size and event probability follows a logarithmic trend, as predicted if pupil signals self-information. To step in this direction, the data from aforementioned studies is plotted against quantified surprise values in Fig. 1 (see squares in figure). This analysis is restricted to studies that reported probabilities quantitatively and measured pupil size in millimetres or percents. Precise comparison across studies is not possible given that detailed conditions are not available (i.e. time and performance pressure, lighting conditions, baseline arousal levels, etc.) and that measurement methods may differ. However, this visualization already suggests that pupil dilation is linearly proportional to self-information, within and across studies.

**Figure 1.**
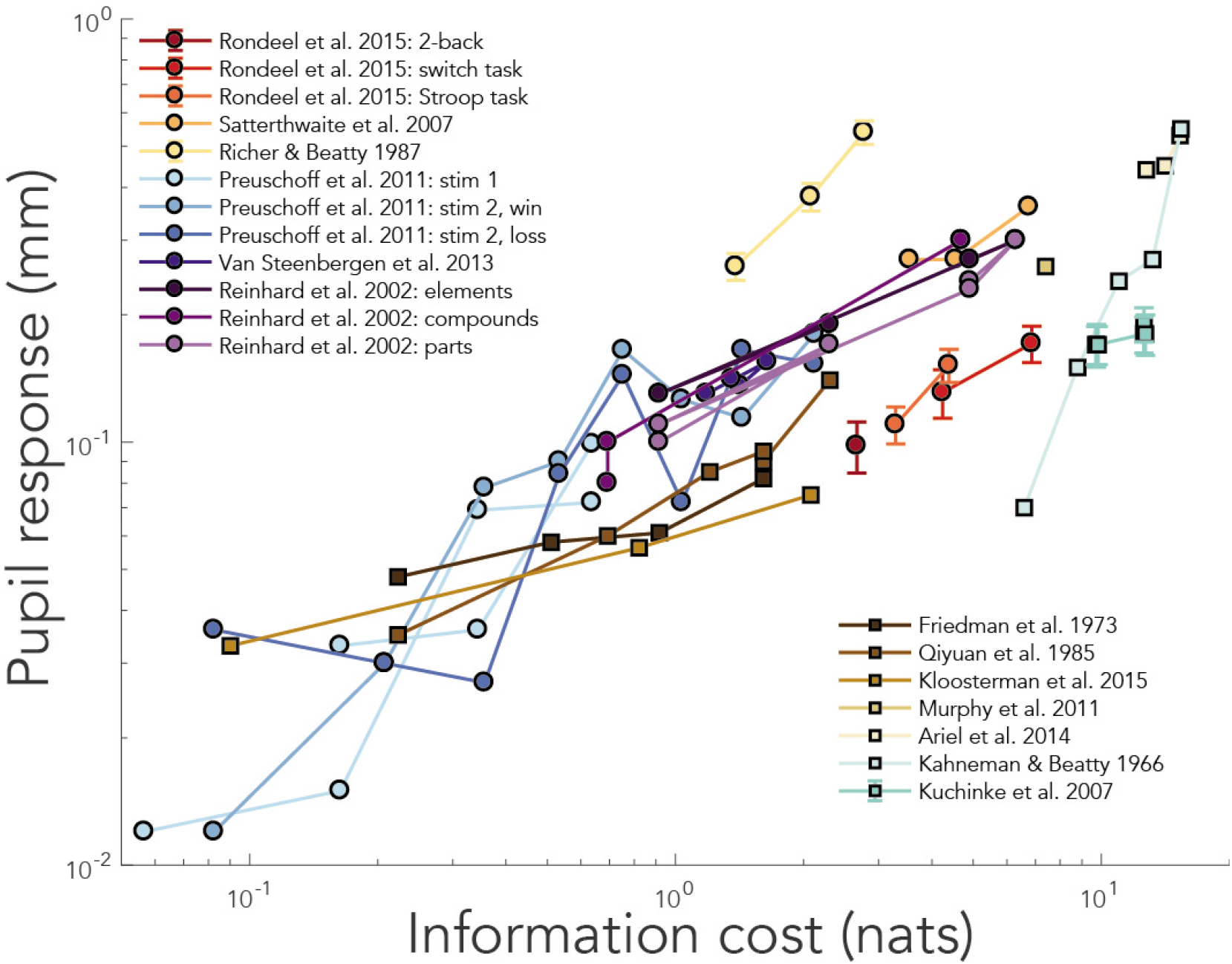
Relationship between information cost and pupil dilation in previous studies. Information cost was quantified as the KL divergence between prior and posterior beliefs. Squares in the graph illustrate pupillary responses to discrimination or detection tasks, in which KL divergence simplifies to stimulus self-information. Circles illustrate pupil dilations in response to task variables and decision making. See supplementary information for details.

## Information about task variables

The examples mentioned so far show that *pupil size dilates in proportion to the amount of information needed to encode sensory stimuli*. When a surprising stimulus is presented, self-information is large and pupils dilate. However, sensory stimuli such as cues, can also carry information about other, separate events. Pupillary response to such cases was investigated in Preuschoff et al. [7], in which stimuli informed participants on their winning probability. Subjects had to bet on which of two cards, whose values were revealed afterwards, was going to be larger. In this study, Preuschoff and colleagues looked at the pupil response to the display of the first card value. Here all values (from 1 to 10) were equally likely, such that self-information was equal in all conditions. However, some cards provided more information than others about the chance of having a winning or losing bet. For example, when the first card was a 10, there was a guarantee of winning/losing if participant had bet on the first card being larger/smaller (there were no ties in the game). Conversely, a 5 provided little information about the chance of winning, since probabilities were still close to 50-50. Such gradual gain of information about the probability distribution of a variable (chance of winning in the present case) can be quantified by the Kullback-Leibler (KL) divergence between *prior* and *posterior* variable distributions. KL divergence can be interpreted as the amount of information gained about the true probability distribution of a variable, after receiving new data. KL divergence provides a generalized measure of information gain that is equivalent to self-information in the case of detection or discrimination tasks. Remarkably, the pupillary response to first card value presentation in Preuschoff et al. [7] followed closely the KL divergence between subjects’ belief on winning probability before and after observing the first card value (see light blue circles in figure 1 and panel A in figure 2), even though these results were not discussed as such in the paper.

**Figure 2.**
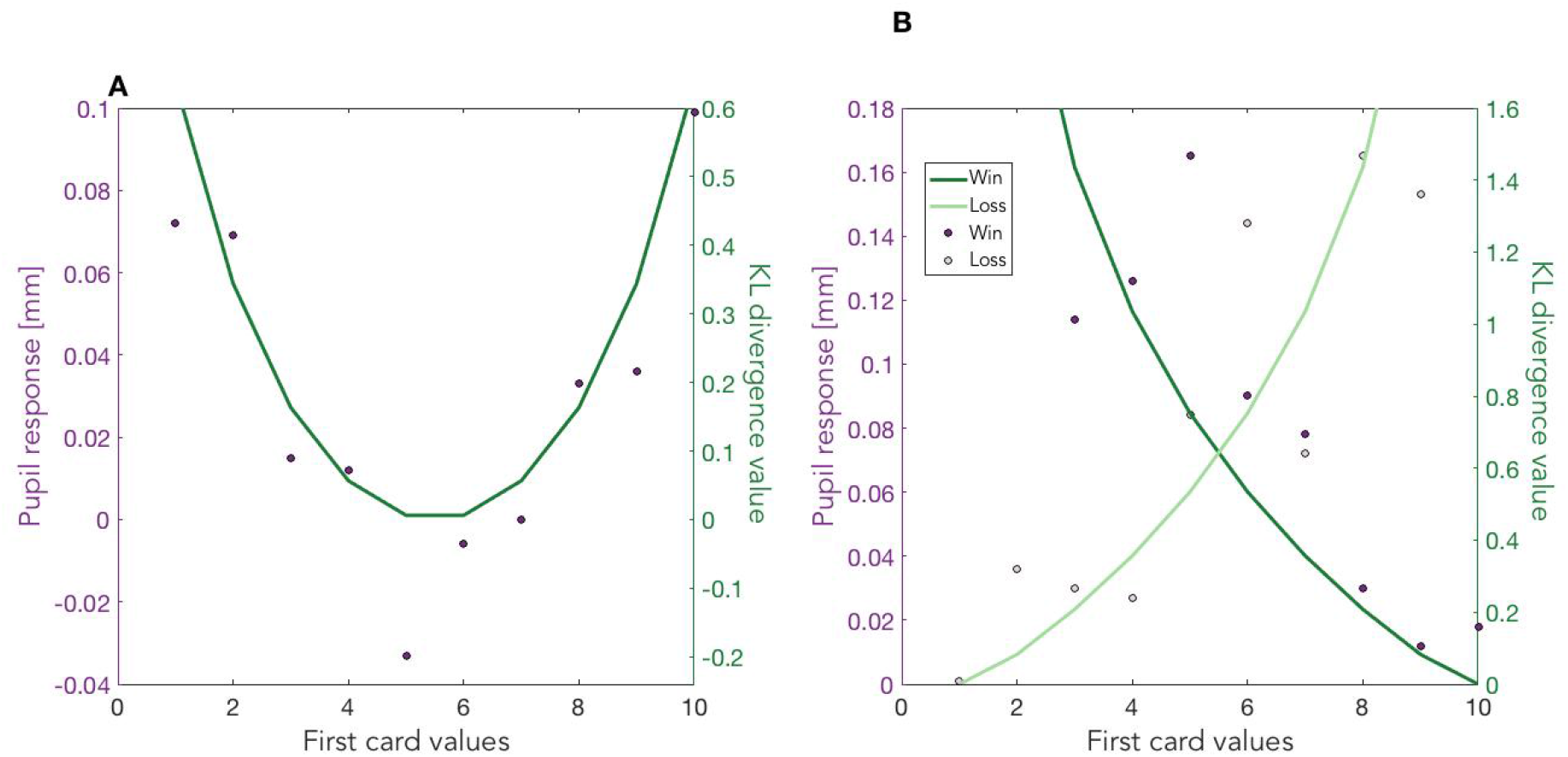
Data from Preuschoff et al. [7] (left y-axis), together with simulations based on KL divergence between probability distribution of winning before and after viewing the stimuli (right y-axis). Responses to first card presentation is shown in panel A, whereas panel B illustrates responses to second card presentation. See supplementary material for details.

When the second card was presented, different situations could occur. The predictions could be confirmed, in which case little information would be gained (e.g. first card was 8, predicting first card being larger, and second card was 5, confirming predictions), or they could be contradicted, in which case a lot of information would be gained (e.g. first card was 8 but second card was 9). Here again, pupil responded in proportion to the amount of information being gained about winning probability, quantified as KL divergence (see figure 1 and panel B in figure 2). The findings of Preuschoff et al. [7] are compelling for several reasons. First, pupil size variations occurred following participants’ choice and were thereby not affected by decision processes or motor responses, reflecting purely inferential processes. Second, they allow us to make clear quantitative predictions in terms of information processing and these predictions are strikingly confirmed.

One difference between surprise and KL divergence models of pupil response is that, *if pupil responded only to surprise, it would always depend on the frequency of occurrence of presented stimulus, independently of task*. In contrast, KL divergence models predict that pupil will respond to the amount of information provided by stimuli about task variables. This difference was exploited in two studies by Reinhard and colleagues [8, 54] in which stimulus probabilities were manipulated in a GO/NOGO tasks. In accordance with the information model, Reinhard et al. showed that pupillary response depended only on the probability of occurrence of the features of the GO/NOGO stimuli that were informative about the task (e.g. when GO was defined by the occurrence of 1-letter as opposed to 2-letter stimuli, the identity of the letter being presented was irrelevant and failed to affect pupil response; see simulated results in figure 1). More generally, several studies have found that pupillary responses to stimuli depend on whether they are attended to or not [31, 32, 34, 55, 56] and that these responses scale with the subjective salience of the stimuli [35, 56, 57]. In attentional blink experiments, targets that follow closely previous target occurrences remain sometimes undetected. In these cases, pupillary response to target occurrence is greatly diminished [32]. Larger pupil dilation is associated with larger distractor interference [58], and increased processing of subliminal cues [59], in agreement with the view that pupil response scales with the quantity of visual information being processed.

## Decision making

When decisions are made in the absence of uncertainty, such as in simple stimulus-response association tasks, the relationship between pupil response and information gain is straightforward. For example, in Richer and colleagues, both reaction time and pupil dilation were shown to vary as a function of the number of stimulus-response associations [38], in accordance with the classical Hick-Hyman law [60]. Here the information cost of the decision can be quantified as the log of the number of possible stimulus-response associations in the task, which is equivalent to the KL divergence between prior and posterior beliefs [61] (see figure 1, yellow circles).

In conditions of uncertainty, the situation is slightly more complex. Satterthwaite and colleagues tested participants on a task similar to that of Preuschoff et al. [7], except that the decision followed, rather than preceded, the display of the first card value [37]. Participants had to pick either the face-up or face-down deck of cards. The second card value was then revealed and the trial was won if the card from the chosen deck was the largest [37]. Interestingly, in that case, the results were exactly opposite those of Preuschoff: when the first card was less informative (e.g. 5), making it more difficult for the subject to choose which deck to pick, the pupil response was larger than when the first number was either small or large, a case for which decision was easier to make. The reaction time associated with the decision followed the same pattern, being larger for less informative values. This observed relationship between reaction time and pupillary dilation has been found in many studies [46, 19, 62, 24, 63, 2, 64] and pupillary responses are best modelled by means of regressors that extend during the whole reaction time period of the trial rather than by brief pulses limited to stimulus onset [18, 22]. These findings suggest that the process from which pupillary dilation originates is maintained during the whole decision process.

The finding that uncertain or conflictual decisions are slower than decisions for which more information is available from stimulus is classical in the decision making literature. It can be modeled as a drift diffusion process in which noisy evidence accumulates until a threshold is reached and in which the rate of accumulation depends on how close the option values are to each other [65, 66]. Drift diffusion models can also be interpreted as time-resolved Bayesian decision making processes in which each accumulation step corresponds to the update of prior to posterior belief [67]. The noisier the evidence, the more updates will tend to go in the wrong direction. Therefore, the summed quantity of information accumulated over the whole decision process is larger when evidence is noisy than when it is not. Thus, results from Satterthwaite et al. [37] can be accounted for by considering the sum of the KL divergences resulting from every update along the drift diffusion process (see figure 3 and light orange circles in figure 1). In Urai et al. [19] and Colizoli et al. [24], pupil size was measured during motion discrimination tasks and was shown to vary in parallel with decision uncertainty and reaction time: it decreased with stimulus strength for correct trials (low uncertainty), but increased with stimulus strength in error trials (high uncertainty). This pattern of results can also be explained by recurring to drift diffusion models of decision making and by assuming variable drift rates [66]. Along the same line, Cheadle et al. [45] showed that during a task in which evidence accumulated over eight successive stimulus presentation, pupillary responses were proportional to the amount of evidence provided by each stimulus. Moreover, this response was modulated by recency and confirmation biases, which both also affected decisions. So pupil responses tracked decision updates, as predicted by our proposal. In de Gee et al. [18] and de Gee et al. [22], pupil responses in detection and 2-alternative forced choice tasks were shown to be inversely proportional to the probability of the choice and hence to the KL divergence between prior and posterior: in conservative participants (biased towards NO), YES choices led to larger responses, while the opposite tended to be found in more liberal participants (biased towards YES). Pupil responses were also shown to vary as a function of the influence of the prior on perceptual decisions in de Gee et al. [22] and Krishnamurthy et al. [21]: when prior beliefs have less weight (because of better control or attentional allocation or because of low prior reliability), more information is extracted from the sensory stimulus, KL divergence is larger and pupil dilates more. Along the same line, when the occurrence of surprising outcomes suggests the task structure may have changed, pupil dilations is even larger [10, 21, 26]. This is because such environmental volatility is associated with increased learning rate and thus increased influence of sensory evidence on internal models of the task. Indeed, the extent to which volatility affected learning rate correlates with the magnitude of the pupil response [10, 21]. Together, these findings on pupillary response to volatility and surprise confirm that pupil diameter scales with how much novel sensory evidence is used to update current belief states.

**Figure 3.**
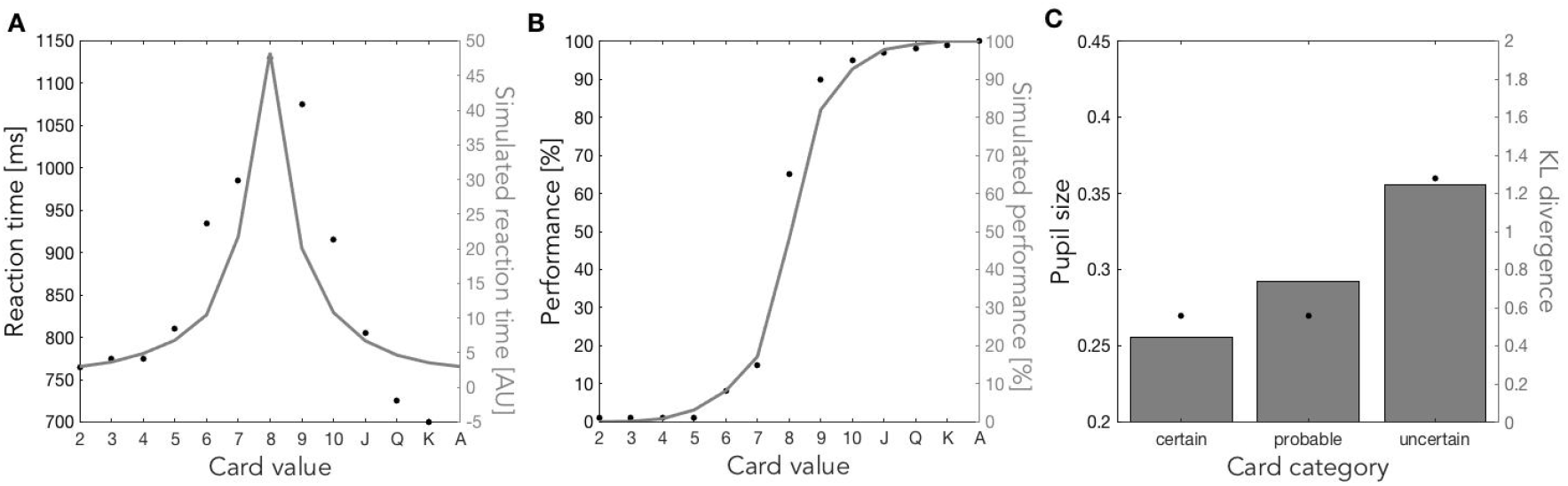
Simulation of reaction times (panel A) and percent correct responses (panel B) from Satterthwaite et al. 2007 by means of a DDM process. Panel C illustrates the resulting KL divergences (grey bars), which follow the same trend (increasing with uncertainty) as the pupil size reported in the original study (black dots). It is noteworthy that the model used to simulate these data has decision threshold as single degree of freedom. See supplementary material for more details.

## Mental effort

Another common findings in the literature is that pupil size varies as a function of task demands and subject’s engagement in the task, suggesting the view that pupillary dilation indexes mental effort [2, 4, 5, 68, 69, 70]. We have recently proposed that mental effort too can be quantified as the average KL divergence between prior and posterior beliefs [61]. Effortful tasks often include large number of associations between stimuli and responses, resulting in low prior beliefs for each association and requiring large updates in order to reach precise posterior beliefs (e.g. N-back task; see simulations of N-back task from Rondeel et al. [68] in Fig. 1, red circle). Other cases of difficult tasks are those in which prior beliefs do not match task statistics (e.g. Stroop task), or in which task statistics change constantly (e.g. switch tasks), also implying large updates and large information costs (see simulations of Stroop and switch tasks from Rondeel et al. [68] in Fig. 1, orange and yellow circle). So the present proposal that pupil size scales with information gain can also be applied to complex tasks and accounts for the classical relation between mental effort and pupillary dilation.

## Tonic pupil size

So far we have restricted our discussion to phasic pupil responses, i.e. the change in pupil size that follows event onset. However, the tonic variations in pupillary diameter, usually measured during baseline epochs that precede trial onsets have also some interesting properties. These tonic pupillary changes have been related to the modes of discharge observed in noradrenergic neurons [29, 63, 48, 47, 50, 30]. Large phasic responses occur when baseline firing rates of noradrenergic neurons are low and would correspond to small tonic pupil size, whereas large baseline noradrenergic activity would be associated with large tonic pupil size but small phasic responses [71, 29, 63, 72, 47]. Indeed, negative correlations between *spontaneous* changes in tonic and phasic pupil size have been reported repeatedly [29, 20, 63, 44, 73, 18, 74, 72], even though *task-induced* or interindividual changes in tonic and phasic pupil size go often in the same direction [12, 21, 10, 30, 25, 75].

The relation between *spontaneous* changes in tonic pupil size and behaviour follows an inverted u-shape, with optimal performance being associated with intermediate pupil size, evoking Yerkes-Dodson law [63, 72, 48, 47, 58]. Large tonic pupil sizes are concurrent with mind-wandering, distractibility and exploratory behaviour [33, 30, 76, 29, 77] while very low tonic pupil sizes are associated with low vigilance and sleepiness [78, 79, 80, 63, 36, 71, 72, 47]. However, in contrast with aforementioned spontaneous changes, increases in tonic size that are *task-induced* occur, on the contrary, in conditions of high task demand: when taxing working memory [81], when counting stimuli silently [82], following changes of contingency [25, 83, 21, 10, 30, 25, 26] or in conditions of high uncertainty [12].

Assuming that tonic variations of pupil size, like phasic task-induced changes, reflect quantitatively the amount of information being processed by the brain may help reconcile these contradictory findings in a parsimonious way. When information is attached to abrupt sensory signal, it leads to phasic dilation whose magnitude is proportional to the KL divergence between prior and posterior beliefs. In the absence of clear onset, tonic pupil size reflects information processing from memory, i.e. manipulation of working memory, planning, mind-wandering, mental imagery or offline learning. Therefore, tonic pupil size would increase when cognitive activity occurs out of sync with task events [76], hence decreasing limited cognitive resources available for main task [61], leading to distractibility and exploratory behaviour, but it would also increase during demanding covert computations on working memory [81, 82, 25]. However, confirming this hypothesis requires quantifying out-of-sync information processing in terms of KL divergence, like we did for phasic pupillary responses. Since we cannot provide such quantified predictions on the basis of current literature, this will have to rely on future experimental studies.

## Relation to alternative theories

Pupillary responses, because of their relation to the noradrenergic system [71], have previously been linked to unexpected uncertainty [27, 84], sometimes taken as synonym to surprise [7, 28, 6] and sometimes as an equivalent of volatility, i.e. how likely the environment dynamics is to change [84, 27, 21, 85, 86, 87]. These two definitions are strongly related since surprising observations suggest that the statistical structure of the environment may have changed [88]. While surprise is event-related and could be linked to phasic pupillary changes [28], volatility varies slowly and could be related to tonic pupil size [27]. Unexpected uncertainty relates also strongly to the problem of exploitation/exploration trade-off, another concept linked to pupillary responses [30, 29, 83, 89]: when confidence in the internal model of the environment drops following surprising observations, exploitation strategies lose value with respect to alternative exploration strategies [84]. However, recent data has shown that variations of tonic pupil size are not indicative of unexpected uncertainty, but are rather a signature of reducible uncertainty (ambiguity resulting from poor model of environment, caused by undersampling; Krishnamurthy et al. [21]) or expected uncertainty (related to the variance of the task; De Berker et al. [12]). This is also in line with the finding that pupil size does not depend only on noradrenaline but also on other neuromediators such as acetylcholine [50], whose function has been associated with encoding of expected uncertainty [27]. Phasic pupillary responses, on the contrary, were shown to correlate with unexpected uncertainty [21]. However, since volatility is a slow-changing property of the environment, this observed correlation with phasic pupillary changes must reflect the fact that, when prior knowledge on environment is unreliable (i.e. volatility is high), more weight is given to new sensory evidence, as opposed to prior biases [90, 84, 27], and model updates between prior and posterior beliefs are more expansive [90], leading to larger pupillary dilations. Overall, current evidence does not seem to favour the view that pupil dilation would be indicative of specific types of uncertainty but, as I argue in the present work, would rather signal information processing, which itself depends strongly on uncertainty conditions.

## Limitations

Notably, two studies reported results that appear to be in contradiction with our information model. In O’Reilly et al. [13], the onset of unexpected saccadic targets led to pupillary dilations, but when these violations of expectation indicated the need to update the internal model of saccade target distributions, pupillary responses were smaller than when these unexpected events were identified as being outliers (identified by their colour). In Van Slooten et al. [23], pupillary response to the outcome of subjects’ choices in a 2-arm bandit task was shown not to depend on modelled expectations: when subjects were thought to expect a large reward, their pupillary response was similar regardless of feedback. Further, the magnitude of the decision-related response scaled with the difference between the available options, and feedback pupillary response was inversely proportional to the model learning rate, both results being in apparent contradiction with previous literature [10, 37, 19] and the present proposal. In both aforementioned cases, pupillary responses were compared to variables of computational models fitted to behaviour, as opposed to direct task variables. These behavioural models are based on assumptions and conclusions drawn from the models are valid only to the extent that these assumptions are justified. For example, in O’Reilly et al. [13] the model assumed participants did not update their internal model when faced with outlier stimuli. However, it could be argued that participants always updated their internal models in the face of surprising targets but had to put extra work to cancel these updates when figuring out that the target was an outlier. So while the results of O’Reilly et al. [13] and Van Slooten et al. [23] appear to contradict our view and invite us to remain cautious in our conclusions, possible alternative interpretations of their data suggest that more investigations should be conducted to resolve this apparent inconsistency.

## Conclusion

In the present paper, the factors that trigger changes in pupil-linked arousal were discussed under the light of information theoretic framework. The hypothesis that pupil size scales with the amount of information being processed, allowed us to explain a wide range of data, sometimes with quantitative predictions. This view applies both to tonic and phasic pupillary responses, the difference being that phasic responses mark information processing triggered by precise event onset while tonic pupillary changes are not precisely aligned to external events.

Beside the factors that trigger pupillary changes, an equally important issue concerns the computational effects of pupil-linked arousal, and more generally, its functional role in brain computations. This issue goes beyond the scope of the present paper and will be discussed in future work.

## Supporting information

Simulation methods

## Acknowledgements

I wish to thank Oleg Solopchuk, Stefano Ioannucci and Sze-Ying Lam for their constructive comments on an earlier version of the manuscript.

## References

1 Laughlin S.B. 1992. Retinal information capacity and the function of the pupil. Ophthalmic Physiol. Opt., 12, 161–164. (doi: 10.1111/j.1475-1313.1992.tb00281.x).

2 Wel Pv. d and Steenbergen Hv. Pupil dilation as an index of effort in cognitive control tasks: A review, 2018. ISSN 15315320.

3 Beatty J. 1982. Task-evoked pupillary responses, processing load, and the structure of processing resources. Psychol. Bull., 91, 276–292. (doi: 10.1037/0033-2909.91.2.276).

4 Kahneman D and Beatty J. 1966. Pupil diameter and load on memory. Science (80-.)., 154, 1583–1585. (doi: 10.1126/science.154.3756.1583).

5 Hess E. H and Polt J. M. 1964. Pupil size in relation to mental activity during simple problemsolving. Science (80-.)., 143, 1190–1192. (doi: 10.1126/science.143.3611.1190).

6 Lavín C, San Martín R, and Rosales Jubal E. 2014. Pupil dilation signals uncertainty and surprise in a learning gambling task. Front. Behav. Neurosci., 7, 218. (doi: 10.3389/fnbeh.2013.00218).

7 Preuschoff K, ’t Hart B. M, and Einhäuser W. 2011. Pupil dilation signals surprise: Evidence for noradrenaline’s role in decision making. Front. Neurosci., 5, 1–12. (doi: 10.3389/fnins.2011.00115).

8 Reinhard G and Lachnit H. 2002. The effect of stimulus probability on pupillary response as an indicator of cognitive processing in human learning and categorization. Biol. Psychol., 60, 199–215.(doi: 10.1016/S0301-0511(02)00031-5).

9 Kloosterman N. A, Meindertsma T, Loon A. Mv, Lamme V. A, Bonneh Y. S, and Donner T. H. 2015. Pupil size tracks perceptual content and surprise. Eur. J. Neurosci., 41, 1068–1078. (doi: 10.1111/ejn.12859).

10 Nassar M. R, Rumsey K. M, Wilson R. C, Parikh K, Heasly B, and Gold J. I. 2012. Rational regulation of learning dynamics by pupil-linked arousal systems. Nat. Neurosci., 15, 1040–1046. (doi: 10.1038/nn.3130).

11 Nakayama K. 2017. Pupil responses to high-level image content Marnix Naber. J. Vis., 13, 1–8. (doi: 10.1167/13.6.7.doi).

12 De Berker A. O, Rutledge R. B, Mathys C, Marshall L, Cross G. F, Dolan R. J, and Bestmann S. 2016. Computations of uncertainty mediate acute stress responses in humans. Nat. Commun., 7, 10996. (doi: 10.1038/ncomms10996).

13 O’Reilly J. X, Schüffelgen U, Cuell S. F, Behrens T. E. J, Mars R. B, and Rushworth M. F. S. 2013. Dissociable effects of surprise and model update in parietal and anterior cingulate cortex. Proc. Natl. Acad. Sci. U. S. A., 110, E3660–9. (doi: 10.1073/pnas.1305373110).

14 Qiyuan J, Richer F, Wagoner B. L, and Beatty J. 1985. The Pupil and 8 Stimulus Probability. Psychophysiology, 22, 530–534. (doi: 10.1111/j.1469-8986.1985.tb01645.x).

15 Alamia A, VanRullen R, Pasqualotto E, Mouraux A, and Zenon A. 2019. Pupil-linked arousal responds to unconscious surprisal. J. Neurosci., 39, 3010–18. (doi: 10.1523/jneurosci.3010-18.2019).

16 Bradley M. B, Miccoli L. M., Escrig M. a, and Lang P. J. 2008. The pupil as a measure of emotional arousal and automatic activation (Author Manuscript). Psychophysiology, 45, 602. (doi: 10.1111/j.1469-8986.2008.00654.x.The).

17 Cavanagh J. F, Wiecki T. V, Kochar A, and Frank M. J. 2014. Eye tracking and pupillometry are indicators of dissociable latent decision processes. J. Exp. Psychol. Gen., 143, 1476–1488. (doi: 10.1037/a0035813).

18 Gee J. Wd, Knapen T, and Donner T. H. 2014. Decision-related pupil dilation reflects upcoming choice and individual bias. Proc. Natl. Acad. Sci., 111, E618–E625. (doi: 10.1073/pnas.1317557111).

19 Urai A. E, Braun A, and Donner T. H. 2017. Pupil-linked arousal is driven by decision uncertainty and alters serial choice bias. Nat. Commun., 8, 14637. (doi: 10.1038/ncomms14637).

20 Murphy P. R, Vandekerckhove J, and Nieuwenhuis S. 2014. Pupil-Linked Arousal Determines Variability in Perceptual Decision Making. PLoS Comput. Biol., 10, e1003854. (doi: 10.1371/journal.pcbi.1003854).

21 Krishnamurthy K, Nassar M. R, Sarode S, and Gold J. I. 2017. Arousal-related adjustments of perceptual biases optimize perception in dynamic environments. Nat. Hum. Behav., 1, 0107. (doi: 10.1038/s41562-017-0107).

22 Gee J. Wd, Colizoli O, Kloosterman N. A, Knapen T, Nieuwenhuis S,and Donner T. H. 2017. Dynamic modulation of decision biases by brainstem arousal systems. Elife, 6, 1–36. (doi: 10.7554/elife.23232).

23 Van Slooten J. C, Jahfari S, Knapen T, and Theeuwes J. 2018. How pupil responses track value-based decision-making during and after reinforcement learning. PLoS Comput. Biol., 14, e1006632. (doi: 10.1371/journal.pcbi.1006632).

24 Colizoli O, Gee J. Wd, Urai A. E, and Donner T. H. 2018. Task-evoked pupil responses reflect internal belief states. Sci. Rep., 8, 13702. (doi: 10.1038/s41598-018-31985-3).

25 Van Slooten J. C, Jahfari S, Knapen T, and Theeuwes J. 2017. Individual differences in eye blink rate predict both transient and tonic pupil responses during reversal learning. PLoS One, 12, 1–20. (doi: 10.1371/journal.pone.0185665).

26 Browning M, Behrens T. E, Jocham G, O’Reilly J. X, and Bishop S. J. 2015. Anxious individuals have difficulty learning the causal statistics of aversive environments. Nat. Neurosci., 18, 590–596. (doi: 10.1038/nn.3961).

27 Yu A. J and Dayan P. 2005. Uncertainty, neuromodulation, and attention. Neuron, 46, 681–692. (doi: 10.1016/j.neuron.2005.04.026).

28 Dayan P and Yu A. J. 2006. Phasic norepinephrine: A neural interrupt signal for unexpected events. Netw. Comput. Neural Syst., 17, 335–350. (doi: 10.1080/09548980601004024).

29 Gilzenrat M. S, Nieuwenhuis S, Jepma M, and Cohen J. D. 2010. Pupil diameter tracks changes in control state predicted by the adaptive gain theory of locus coeruleus function. Cogn. Affect. Behav. Neurosci., 10, 252–269. (doi: 10.3758/CABN.10.2.252).

30 Jepma M and Nieuwenhuis S. 2011. Pupil diameter predicts changes in the exploration-exploitation trade-off: Evidence for the adaptive gain theory. J. Cogn. Neurosci., 23, 1587–1596. (doi: 10.1162/jocn.2010.21548).

31 Kang O. E, Huffer K. E, and Wheatley T. P. 2014. Pupil dilation dynamics track attention to high-level information. PLoS One, 9, e102463. (doi: 10.1371/journal.pone.0102463).

32 Wierda S. M, Rijn Hv, Taatgen N. A, and Martens S. 2012. Pupil dilation deconvolution reveals the dynamics of attention at high temporal resolution. Proc. Natl. Acad. Sci., 109, 8456–8460. (doi: 10.1073/pnas.1201858109).

33 Smallwood J, Brown K. S, Tipper C, Giesbrecht B, Franklin M. S, Mrazek M. D, Carlson J. M, and Schooler J. W. 2011. Pupillometric evidence for the decoupling of attention from perceptual input during offline thought. PLoS One, 6, e18298. (doi: 10.1371/journal.pone.0018298).

34 Liao H. I, Yoneya M, Kidani S, Kashino M, and Furukawa S. 2016. Human pupillary dilation response to deviant auditory stimuli: Effects of stimulus properties and voluntary attention. 10,. (doi: 10.3389/fnins.2016.00043).

35 Liao H. I, Kidani S, Yoneya M, Kashino M, and Furukawa S. 2016. Correspondences among pupillary dilation response, subjective salience of sounds, and loudness. Psychon. Bull. Rev., 23, 412–425. (doi:10.3758/s13423-015-0898-0).

36 Van Den Brink R. L, Murphy P. R, and Nieuwenhuis S. 2016. Pupil diameter tracks lapses of attention. PLoS One, 11, e0165274. (doi: 10.1371/journal.pone.0165274).

37 Satterthwaite T. D, Green L, Myerson J, Parker J, Ramaratnam M, and Buckner R. L. 2007. Dissociable but inter-related systems of cognitive control and reward during decision making: Evidence from pupillometry and event-related fMRI. Neuroimage, 37, 1017–1031. (doi: 10.1016/j.neuroimage.2007.04.066).

38 Richer F and Beatty J. 1987. Contrasting Effects of Response Uncertainty on the Task-Evoked Pupillary Response and Reaction Time. Psychophysiology, 24, 258–262. (doi:10.1111/j.1469-8986.1987.tb00291.x).

39 Lempert K. M, Chen Y. L, and Fleming S. M. 2015. Relating pupil dilation and metacognitive confidence during auditory decision-making. PLoS One, 10, e0165274. (doi: 10.1371/journal.pone.0126588).

40 Manohar S. G, Finzi R. D, Drew D, and Husain M. 2017. Distinct Motivational Effects of Contingent and Noncontingent Rewards. Psychol. Sci., 28, 1016–1026. (doi: 10.1177/0956797617693326).

41 Jepma M, Murphy P. R, Nassar M. R, Rangel-Gomez M, Meeter M, and Nieuwenhuis S. 2016. Catecholaminergic Regulation of Learning Rate in a Dynamic Environment. PLoS Comput. Biol., 12, e1005171. (doi: 10.1371/journal.pcbi.1005171).

42 McGuire J. T, Nassar M. R, Gold J. I, and Kable J. W. 2014. Functionally Dissociable Influences on Learning Rate in a Dynamic Environment. Neuron, 84, 870–881. (doi: 10.1016/j.neuron.2014.10.013).

43 Murphy P. R, Van Moort M. L, and Nieuwenhuis S. 2016. The pupillary orienting response predicts adaptive behavioral adjustment after errors. PLoS One, 11, e0151763. (doi: 10.1371/journal.pone.0151763).

44 Eldar E, Cohen J. D, and Niv Y. 2013. The effects of neural gain on attention and learning. Nat. Neurosci., 16, 1146–1153. (doi: 10.1038/nn.3428).

45 Cheadle S, Wyart V, Tsetsos K, Myers N, DeGardelle V, HerceCastañón S, and Summerfield C. 2014. Adaptive gain control during human perceptual choice. Neuron, 81, 1429–1441. (doi: 10.1016/j.neuron.2014.01.020).

46 Murphy P. R, Boonstra E, and Nieuwenhuis S. 2016. Global gain modulation generates time-dependent urgency during perceptual choice in humans. Nat. Commun., 7, 14299. (doi: 10.1038/ncomms13526).

47 McGinley M. J, Vinck M, Reimer J, Batista-Brito R, Zagha E, Cadwell C. R, Tolias A. S, Cardin J. A, and McCormick D. A. 2015. Waking State: Rapid Variations Modulate Neural and Behavioral Responses. Neuron, 87, 1143–1161. (doi: 10.1016/j.neuron.2015.09.012).

48 McGinley M. J, David S. V, and McCormick D. A. 2015. Cortical Membrane Potential Signature of Optimal States for Sensory Signal Detection. Neuron, 87, 179–192. (doi: 10.1016/j.neuron.2015.05.038).

49 Reimer J, Froudarakis E, Cadwell C. R, Yatsenko D, Denfield G. H, and Tolias A. S. 2014. Pupil Fluctuations Track Fast Switching of Cortical States during Quiet Wakefulness. Neuron, 84, 355–362. (doi: 10.1016/j.neuron.2014.09.033).

50 Reimer J, McGinley M. J, Liu Y, Rodenkirch C, Wang Q, McCormick D. A, and Tolias A. S. 2016. Pupil fluctuations track rapid changes in adrenergic and cholinergic activity in cortex. Nat. Commun., 7, 13289. (doi: 10.1038/ncomms13289).

51 Friedman D, Hakerem G, Sutton S, and Fleiss J. L. 1973. Effect of stimulus uncertainty on the pupillary dilation response and the vertex evoked potential. Electroencephalogr. Clin. Neurophysiol., 34, 475–484. (doi: 10.1016/0013-4694(73)90065-5).

52 Kuchinke L, Võ M. L, Hofmann M, and Jacobs A. M. 2007. Pupillary responses during lexical decisions vary with word frequency but not emotional valence. Int. J. Psychophysiol., 65, 132–140. (doi: 10.1016/j.ijpsycho.2007.04.004).

53 Braem S, Coenen E, Bombeke K, Bochove M. Ev, and Notebaert W. 2015. Open your eyes for prediction errors. Cogn. Affect. Behav. Neurosci., 15, 374–380. (doi: 10.3758/s13415-014-0333-4).

54 Reinhard G, Lachnit H, and König S. 2007. Effects of stimulus probability on pupillary dilation and reaction time in categorization. Psychophysiology, 44, 469–475. (doi: 10.1111/j.1469-8986.2007.00512.x).

55 Kang O and Wheatley T. 2015. Pupil dilation patterns reflect the contents of consciousness. Conscious. Cogn., 35, 128–135. (doi: 10.1016/j.concog.2015.05.001).

56 Ariel R and Castel A. D. 2014. Eyes wide open: Enhanced pupil dilation when selectively studying important information. Exp. Brain Res., 232, 337–344. (doi: 10.1007/s00221-013-3744-5).

57 Damsma A and Rijn Hv. 2017. Pupillary response indexes the metrical hierarchy of unattended rhythmic violations. Brain Cogn., 111, 95–103. (doi: 10.1016/j.bandc.2016.10.004).

58 Ebitz R. B, Pearson J. M, and Platt M. L. 2014. Pupil size and social vigilance in rhesus macaques. Front. Neurosci.,8, 100. (doi: 10.3389/fnins.2014.00100).

59 Chiesa P. A, Liuzza M. T, Acciarino A, and Aglioti S. M. 2015. Subliminal perception of others’ physical pain and pleasure. Exp. Brain Res., 233, 2373–2382. (doi: 10.1007/s00221-015-4307-8).

60 Hick W. E. 1952. On the Rate of Gain of Information. Q. J. Exp. Psychol., 4, 11–26. (doi: 10.1080/17470215208416600).

61 Zénon A, Solopchuk O, and Pezzulo G. 2018. An information-theoretic perspective on the costs of cognition. Neuropsychologia, 123, 5–18. (doi:10.1016/j.neuropsychologia.2018.09.013).

62 Gomes C. A, Montaldi D, and Mayes A. 2015. The pupil as an indicator of unconscious memory: Introducing the pupil priming effect. Psychophysiology, 52, 754–769. (doi: 10.1111/psyp.12412).

63 Murphy P. R, Robertson I. H, Balsters J. H, and O’connell R. G. 2011. Pupillometry and P3 index the locus coeruleus-noradrenergic arousal function in humans. Psychophysiology, 48, 1532–1543. (doi: 10.1111/j.1469-8986.2011.01226.x).

64 Steenbergen Hv and Band G. P. H. 2013. Pupil dilation in the Simon task as a marker of conflict processing. Front. Hum. Neurosci., 7, 215. (doi: 10.3389/fnhum.2013.00215).

65 Pedersen M. L, Frank M. J, and Biele G. 2017. The drift diffusion model as the choice rule in reinforcement learning. Psychon. Bull. Rev., 24, 1234–1251. (doi: 10.3758/s13423-016-1199-y).

66 Ratcliff R and McKoon G. 2008. The diffusion decision model: Theory and data for two-choice decision tasks. Neural Comput., 20, 873–922. (doi:10.1162/neco.2008.12-06-420).

67 Bitzer S, Park H, Blankenburg F, and Kiebel S. J. 2014. Perceptual decision making:drift-diffusion model is equivalent to a Bayesian model. Front. Hum. Neurosci., 8, 102. (doi: 10.3389/fnhum.2014.00102).

68 Rondeel E. W. M, Steenbergen Hv, Holland R. W, and Knippenberg Av. 2015. A closer look at cognitive control: differences in resource allocation during updating, inhibition and switching as revealed by pupillometry. Front. Hum. Neurosci., 9, 494. (doi: 10.3389/fnhum.2015.00494).

69 Bijleveld E. 2018. The feeling of effort during mental activity. Conscious. Cogn., 63, 218–227. (doi: 10.1016/j.concog.2018.05.013).

70 Mathôt S. 2018. Pupillometry: Psychology, Physiology, and Function. J. Cogn., 1, 16. (doi: 10.5334/joc.18).

71 Aston-Jones G and Cohen J. D. 2005. An integrative theory of locus coeruleus-norepinephrine function: adaptive gain and optimal performance. Annu. Rev. Neurosci., 28, 403–50. (doi: 10.1146/annurev.neuro.28.061604.135709).

72 Kempen Jv, Loughnane G. M, Newman D. P, Kelly S. P, Thiele A, O’Connell R. G, and Bellgrove M. A. 2019. Behavioural and neural signatures of perceptual decision-making are modulated by pupil-linked arousal. Elife, 8, 1–43. (doi: 10.7554/elife.42541).

73 Joshi S, Li Y, Kalwani R. M, and Gold J. I. 2016. Relationships between Pupil Diameter and Neuronal Activity in the Locus Coeruleus, Colliculi, and Cingulate Cortex. Neuron, 89, 221–34. (doi: 10.1016/j.neuron.2015.11.028).

74 Knapen T, De Gee J. W, Brascamp J, Nuiten S, Hoppenbrouwers S, and Theeuwes J. 2016. Cognitive and ocular factors jointly determine pupil responses under equiluminance. PLoS One, 11, e0155574. (doi: 10.1371/journal.pone.0155574).

75 Van Der Meer E, Beyer R, Horn J, Foth M, Bornemann B, Ries J, Kramer J, Warmuth E, Heekeren H. R, and Wartenburger I. 2010. Resource allocation and fluid intelligence: Insights from pupillometry. Psychophysiology, 47, 158–169. (doi: 10.1111/j.1469-8986.2009.00884.x).

76 Ebitz R. B and Platt M. L. 2015. Neuronal activity in primate dorsal anterior cingulate cortex signals task conflict and predicts adjustments in pupil-linked arousal. Neuron, 85, 628–640. (doi: 10.1016/j.neuron.2014.12.053).

77 Bijleveld E, Custers R, and Aarts H. 2009. The unconscious eye opener: Pupil dilation reveals strategic recruitment of resources upon presentation of subliminal reward cues. Psychol. Sci., 20, 1313–1315. (doi: 10.1111/j.1467-9280.2009.02443.x).

78 Wilhelm B, Giedke H, Lüdtke H, Bittner E, Hofmann A, and Wilhelm H. 2001. Daytime variations in central nervous system activation measured by a pupillographic sleepiness test. J. Sleep Res., 10, 1–7. (doi: 10.1046/j.1365-2869.2001.00239.x).

79 Kristjansson S. D, Stern J. A, Brown T. B, and Rohrbaugh J. W. 2009. Detecting phasic lapses in alertness using pupillometric measures. Appl. Ergon., 40, 978–986. (doi: 10.1016/j.apergo.2009.04.007).

80 Nishiyama J, Tanida K, Kusumi M, and Hirata Y. 2007. The pupil as a possible premonitor of drowsiness. Annu. Int. Conf. IEEE Eng. Med. Biol. – Proc., 2007, 1586–1589. (doi: 10.1109/IEMBS.2007.4352608).

81 Peysakhovich V, Vachon F, and Dehais F. 2017. The impact of luminance on tonic and phasic pupillary responses to sustained cognitive load. Int. J. Psychophysiol., 112, 40–45. (doi: 10.1016/j.ijpsycho.2016.12.003).

82 Steiner G. Z and Barry R. J. 2011. Pupillary responses and event-related potentials as indices of the orienting reflex. Psychophysiology, 48, 1648–1655. (doi:10.1111/j.1469-8986.2011.01271.x).

83 Pajkossy P, Szőllősi Á, Demeter G, and Racsmány M. 2017. Tonic noradrenergic activity modulates explorative behavior and attentional set shifting: Evidence from pupillometry and gaze pattern analysis. Psychophysiology, 54, 1839–1854. (doi: 10.1111/psyp.12964).

84 Parr T and Friston K. J. 2017. Uncertainty, epistemics and active Inference. J. R. Soc. Interface, 14, 20170376. (doi: 10.1098/rsif.2017.0376).

85 Yu A and Dayan P. 2003. Expected and unexpected uncertainty: ACh and NE in the neocortex. Nips 15, 15, 157–164. (doi:citeulike-article-id:496920).

86 Payzan-LeNestour E, Dunne S, Bossaerts P, and O’Doherty 2013. The Neural Representation of Unexpected Uncertainty during Value-Based Decision Making. Neuron, 79, 191–201. (doi: 10.1016/j.neuron.2013.04.037).

87 Heilbron M and Meyniel F. 2019. Confidence resets reveal hierarchical adaptive learning in humans. PLoS Comput. Biol., 15, e1006972. (doi: 10.1371/journal.pcbi.1006972).

88 Soltani A and Izquierdo A. 2019. Adaptive learning under expected and unexpected uncertainty. (doi: 10.1038/s41583-019-0180-y).

89 Lewis G. J and Bates T. C. 2015. Pupil diameter tracks the exploration–Exploitation trade-off during analogical reasoning and explains individual differences in fluid intelligence. J. Cogn. Neurosci., 28, 308–318. (doi: 10.1162/jocn).

90 Moens V and Zénon A. 2019. Learning and forgetting using reinforced Bayesian change detection. PLoS Comput. Biol., 15, e1006713. (doi: 10.1371/journal.pcbi.1006713).

