## Supplementary material for "Eye pupil signals information gain": Simulation methods

### Supplemental information

#### Simulation of past results

**Rondeel, van Steenbergen, Holland, and van Knippenberg (2015): 2-back task.** The average KL divergence between prior and posterior in the 2-back task was quantified as the entropy of item probability distribution (Zénon, Solopchuk, & Pezzulo, 2018). Since probability of letter repetition was 0.378 and assuming equal probabilities among the 25 remaining letters, we obtained an entropy of 2.67: $C = \mathcal{H}(l) = -\sum_{i=1}^{26} p(l_i) \log p(l_i)$, with $p(l_{1-26}) = [0.378, 0.0249, ...0.0249]$. The cost of maintaining past letters in memory was not taken into account, given that this cost should be constant over time and should not affect the pupil response aligned on letter onset.

**Rondeel et al. (2015): switch task and Stroop task.** Computing the cost of Stroop and switching tasks required making some assumption on the strength and flexibility of prior belief on context (Zénon et al., 2018). Assuming context learning rate of 0.3 (like in Zénon et al. (2018)), we found that informational cost of the switching task was 4.25 for no switch trials and 6.85 for switch trials (see Zénon et al. (2018) for details). Finally, the cost of the Stroop task was estimated at 3.24 and 5.54 for congruent and incongruent trials, respectively (assuming prior belief on word naming context of 0.99).

**Friedman, Hakerem, Sutton, and Fleiss (1973).** Information cost for each stimulus was computed as their self-information: $C = -log(p)$, with p being equal to 0.2, 0.4, 0.6 or 0.8.

**Qiyuan, Richer, Wagoner, and Beatty (1985).** Information cost was computed as in Friedman et al. (1973) with p equal to 0.1, 0.2, 0.3, 0.5 or 0.8.

**Satterthwaite et al. (2007).** Information cost was computed as the sum, along all steps of the drift diffusion process, of the KL divergences between prior and posterior beliefs on choice. In each step of the drift diffusion process, the beta prior over the chance of the face down card being the winning card (with $\alpha$ and $\beta$ parameters set to one at the start) is updated by sampling randomly a card and by adding one to $\alpha$ if that card value is superior to the one with the face up, or adding one to $\beta$ in the other case. The KL divergence between prior and posterior is computed in each of these steps. A log-likelihood ratio is also computed as $\log(I_{0.5}(\alpha,\beta)) - \log(1 - I_{0.5}(\alpha,\beta))$, with $I_{0.5}(\alpha,\beta)$ being the cumulative density function of the beta distribution evaluated at p=0.5. The process is interrupted whenever the absolute value of the log-likelihood ratio reaches 3.5. This threshold value is the only free parameter of the model. These simulations were used in Figure 1 and 3.

**Kloosterman et al. (2015).** Information cost in Kloosterman et al. (2015) was computed as the self-information relative to stimulus occurence, based on the hazard rate functions described in the paper. Since pupil size are reported in % in Kloosterman et al. (2015), pupil sizes are converted to mm assuming a baseline pupil size of 3 mm.

**Murphy, Robertson, Balsters, and O'connell (2011).** Similarly to Kloosterman et al. (2015), response to targets were determined as a function of their time of occurrence, based on a hazard rate function with uniform delay distribution. In addition, log(5) was added to the obtained value to account for the 20% chance of target display.

$C = -\sum p \log(h) + log(5)$, with $p = \frac{1}{890}$ for each of the possible delays, equally and linearly spaced between 2 and 2.9 seconds, and $h = \frac{p}{\mathrm{cdf}(p)}$, with cdf the cumulative distribution function.

**Ariel and Castel (2014).** $C = -\log(\frac{7240}{131\times10^6}) - R\log(\frac{7240}{131\times10^6})$, with R corresponding to the recall probability reported in the Results section of Ariel and Castel (2014), to account for the variability of the encoding. The numeric values correspond to reported word frequency.

**Kahneman and Beatty (1966).** $C = m\log(9)$, with m varying from 3 to 7 and representing the number of stimuli to maintain in memory.

**Richer and Beatty (1987).** $C = \log(4) + \log(r)$, with r representing the number of possible responses (Hick, 1952).

**Kuchinke, Võ, Hofmann, and Jacobs (2007).** Word frequency was modeled as a gamma distribution whose mean and standard deviation were equal to the ones reported in the paper (table 1). Cost was then modeled as the entropy of these distributions.

**Preuschoff, 't Hart, and Einhäuser (2011).** Cost for first stimulus was computed as the KL divergence between uniform prior belief and a posterior computed as the probability of winning given the value of the first stimulus. Cost for second stimulus was the KL between the posterior following first stimulus onset and the posterior following second stimulus onset, with p=0.99 for the actual outcome and p=0.01 for the case that did not occur.

**van Steenbergen and Band (2013).** Behavioural effect of switches was accounted by using the same approach as for switch tasks in Rondeel et al. (2015), as described in Zénon et al. (2018). Initial preference for congruent was represented by an assymetric prior joint distribution of stimulus occurrence with p=0.275 for congruent and p=0.225 for incongruent contexts. Similarly as the simulation of Rondeel et al. (2015) above and as Zénon et al. (2018), a learning rate of 0.3 was assumed, and the joint probability was updated following occurrence of congruent or incongruent stimuli. Cost was then computed as the negative logarithm of the resulting probabilities.

**Reinhard and Lachnit (2002).** In Reinhard and Lachnit (2002), the cost of the GO signal was computed on the basis of Fitt's law, as $\log_2(\frac{D}{W} + 1)$, with D the distance = 30cm and W the width of the key = 2cm. Self-information ($-\log p$ with p probability of occurrence) was added to these values.

#### References


Ariel, R., & Castel, A. D. (2014). Eyes wide open: Enhanced pupil dilation when selectively studying important information. *Exp. Brain Res.*, *232*(1), 337–344. doi: 10.1007/s00221-013-3744-5

Friedman, D., Hakerem, G., Sutton, S., & Fleiss, J. L. (1973). Effect of stimulus uncertainty on the pupillary dilation response and the vertex evoked potential. *Electroencephalogr. Clin. Neurophysiol.*, *34*(5), 475–484. doi: 10.1016/0013-4694(73)90065-5

Hick, W. E. (1952). On the Rate of Gain of Information. *Q. J. Exp. Psychol.*, *4*(1), 11–26. Retrieved from `http://journals.sagepub.com/doi/10.1080/17470215208416600` doi: 10.1080/17470215208416600

Kahneman, D., & Beatty, J. (1966). Pupil diameter and load on memory. *Science (80-. ).*, *154*(3756), 1583–1585. doi: 10.1126/science.154.3756.1583

Kloosterman, N. A., Meindertsma, T., van Loon, A. M., Lamme, V. A., Bonneh, Y. S., & Donner, T. H. (2015). Pupil size tracks perceptual content and surprise. *Eur. J. Neurosci.*, *41*(8), 1068–1078. doi: 10.1111/ejn.12859

Kuchinke, L., Võ, M. L., Hofmann, M., & Jacobs, A. M. (2007). Pupillary responses during lexical decisions vary with word frequency but not emotional valence. *Int. J. Psychophysiol.*, *65*(2), 132–140. Retrieved from `http://linkinghub.elsevier.com/retrieve/pii/S0167876007000724 papers2://publication/doi/10.1016/j.ijpsycho.2007.04.004` doi: 10.1016/j.ijpsycho.2007.04.004

Murphy, P. R., Robertson, I. H., Balsters, J. H., & O'connell, R. G. (2011). Pupillometry and P3 index the locus coeruleus-noradrenergic arousal function in humans. *Psychophysiology*, *48*(11), 1532–1543. doi: 10.1111/j.1469-8986.2011.01226.x

Preuschoff, K., 't Hart, B. M., & Einhäuser, W. (2011). Pupil dilation signals surprise: Evidence for noradrenaline's role in decision making. *Front. Neurosci.*, *5*(SEP), 1–12. doi: 10.3389/fnins.2011.00115

Qiyuan, J., Richer, F., Wagoner, B. L., & Beatty, J. (1985). The Pupil and Stimulus Probability. *Psychophysiology*, *22*(5), 530–534. doi: 10.1111/j.1469-8986.1985.tb01645.x

Reinhard, G., & Lachnit, H. (2002). The effect of stimulus probability on pupillary response as an indicator of cognitive processing in human learning and categorization. *Biol. Psychol.*, *60*(2-3), 199–215. doi: 10.1016/S0301-0511(02)00031-5

Richer, F., & Beatty, J. (1987). Contrasting Effects of Response Uncertainty on the Task-Evoked Pupillary Response and Reaction Time. *Psychophysiology*, *24*(3), 258–262. doi: 10.1111/j.1469-8986.1987.tb00291.x

Rondeel, E. W. M., van Steenbergen, H., Holland, R. W., & van Knippenberg, A. (2015). A closer look at cognitive control: differences in resource allocation during updating, inhibition and switching as revealed by pupillometry. *Front. Hum. Neurosci.*, *9*. doi: 10.3389/fnhum.2015.00494

Satterthwaite, T. D., Green, L., Myerson, J., Parker, J., Ramaratnam, M., & Buckner, R. L. (2007). Dissociable but inter-related systems of cognitive control and reward during decision making: Evidence from pupillometry and event-related fMRI. *Neuroimage*, *37*(3), 1017–1031. doi: 10.1016/j.neuroimage.2007.04.066

van Steenbergen, H., & Band, G. P. H. (2013). Pupil dilation in the Simon task as a marker of conflict processing. *Front. Hum. Neurosci.*, *7*. doi: 10.3389/fnhum.2013.00215

Zénon, A., Solopchuk, O., & Pezzulo, G. (2018). An information-theoretic perspective on the costs of cognition. *Neuropsychologia*, *123*, 5–18. doi: 10.1016/j.neuropsychologia.2018.09.013
